# A theoretical framework to study the influence of electrical fields on mesenchymal stem cells

**DOI:** 10.1101/2020.05.04.075697

**Authors:** Jonathan Dawson, Poh Soo Lee, Ursula van Rienen, Revathi Appali

## Abstract

Mesenchymal stem cell dynamics involves cell proliferation and cell differentiation into cells of distinct functional type, such as osteoblasts, adipocytes, or chondrocytes. Electrically active implants influence these dynamics for the regeneration of the cells in damaged tissues. How applied electric field influences processes of individual stem cells is a problem mostly unaddressed. The mathematical approaches to study stem cell dynamics have focused on the stem cell population as a whole, without resolving individual cells and intracellular processes. In this paper, we present a theoretical framework to describe the dynamics of a population of stem cells, taking into account the processes of the individual cells. We study the influence of the applied electric field on the cellular processes. We test our mean-field theory with the experiments from the literature, involving *in vitro* electrical stimulation of stem cells. We show that a simple model can quantitatively describe the experimentally observed time-course behavior of the total number of cells and the total alkaline phosphate activity in a population of mesenchymal stem cells. Our results show that the stem cell differentiation rate is dependent on the applied electrical field, confirming published experimental findings. Moreover, our analysis supports the cell density-dependent proliferation rate. Since the experimental results are averaged over many cells, our theoretical framework presents a robust and sensitive method for determining the effect of applied electric fields at the scale of the individual cell. These results indicate that the electric field stimulation may be effective in promoting bone regeneration by accelerating osteogenic differentiation.

## 1 INTRODUCTION

Human mesenchymal stem cells (hMSCs) possess a unique capability of self-renewal and differentiation into cells of various types of tissues, such as bone, cartilage, and adipose. Thus the hMSCs are the promising cell types for regenerative medicine and tissue engineering. The gene expression levels of an hMSC are known to be the decisive regulators of hMSCs differentiation. These gene expression levels might be influenced by both cell internal cues [De-Leon and Davidson (2007); Ralston (2008)] and external cues [Eyckmans et al. (2012); Najafabadi et al. (2016); Dingal et al. (2014); Eng (2006); Hess et al. (2012b)]. Experimental studies [Mousavi and Hamdy Doweidar (2015)] have shown that the *in vitro* differentiation of hMSC into cells of distinct functional types can be controlled by external factors. Therefore, stem cell differentiation mediated by external factors is a compelling approach that has led to the development of bio-implants, for clinical applications in regenerative medicine.

The applied electric field (EF) is one of the proven external factors known to influence hMSCs dynamics such as migration [Funk (2015); Banks et al. (2015); Ciombor and Aaron (1993); Schemitsch and Kuzyk (2009)], elongation [Rajnicek et al. (2008); Tandon et al. (2009)], proliferation [Sun et al. (2009); Hartig et al. (2000); Kim (2009); Lohmann et al. (2000)], and differentiation [Rohde et al. (2019); Miyamoto et al. (2019); Petecchia et al. (2015); Hess et al. (2012a); Jansen et al. (2010)]. Comparing these studies, it is evident that the results are inconsistent and show the disparity. While several works have demonstrated an increase in proliferation after exposing cells to EF or electromagnetic field (EMF) [Chang et al. (2004); Hartig et al. (2000); Sun et al. (2009); Kim (2009)], others did not detect significant differences or had recorded reduced cell number following EMF exposure [Lohmann et al. (2000); Jansen et al. (2010); Schwartz et al. (2008)]. Similarly, stimulation effects on osteogenic differentiation are also controversial, ranging from no effects [Chang et al. (2004); Lin and Lin (2011)] to a high increase in the expression of bone-related gene markers [Hartig et al. (2000); Jansen et al. (2010); Schwartz et al. (2008)]. Due to the complex parameters and the different experimental approaches used, it is difficult to compare these results among each other. In addition, the choice of stimulation method can also influence cellular behavior.

These methods consist of direct or indirect electrical stimulation of the tissue [Schemitsch and Kuzyk (2009)]. In the direct stimulation method, the electrodes are placed in contact with the targeted tissue. Some of the disadvantages of direct stimulation are the damage caused to tissues by invasive electrodes and the corrosion of the electrodes due to electrochemical processes [Ciombor and Aaron (1993)]. The indirect stimulation method includes capacitive coupling and inductive coupling of electromagnetic fields (EMF). The capacitive coupling is slightly invasive and provides electrical stimulation to the tissue, whereas non-invasive inductive coupling involves both magnetic and electrical stimulation.

To study the stand-alone effects of the EF on the biological tissue, an *in vitro* setup, which is non-invasive and free from the magnetic fields, was necessary. In this context, Hess et al., have developed a novel *in vitro* transformer-like coupling (TC) setup [Hess et al. (2012b)]. This approach enables a non-invasive electrical stimulation of *in-vitro* culture of hMSCs with homogeneous EF in the cell culture chamber. The TC setup exerts pure EFs to the cell culture, with negligible magnetic field strength (see Section 2.1). Thus allowing direct correlation of observed results solely to EF stimulation.

Besides the experimental evaluations, there is a great interest in mathematical modeling and simulation to (i) further gather an in-depth understanding of the cellular mechanism underlying the stem cell response to EMFs, and (ii) to predict optimal stimulation parameters. Fricke [Fricke (1953)] was the first to introduce an empirical equation for the electric potential induced in an ellipsoidal cell in suspension when exposed to an external EF. The first theoretical description (analytical solution of Laplace equation) for the induced potential in a spherical cell in suspension exposed to external EF was given by Schwan [Schwan (1994)] where a spherical shell representing the membrane approximates the cell. This Schwan model treats the cell as a non-conducting membrane subjected to both constant and alternating external EF [Grosse and Schwan (1992)]. Schwan’s theory has been extended by Kotnik et al. [Kotnik et al. (1997)] by considering the conductivity using constant, oscillating, and pulsed EF. Later other geometries such as cylindrical, spheroidal, and ellipsoidal cells suspended in the medium were investigated [Valic et al. (2003); Maswiwat et al. (2008); Gimsa and Wachner (2001a,b)]. To determine the induced EF in the internal membranes of the cells, the cells were modeled as multiple concentric shells [Kotnik and Miklavčič (2006); Vajrala et al. (2008)]. Several techniques were also employed to examine different cells of complex shapes suspended in an electrolyte, for example, Finite Element Models (FEM) [Meny et al. (2007); Sebastián et al. (2004); Ying and Henriquez (2007); Miller and Henriquez (1988)], Transport Lattice Models (TLM) [Gowrishankar and Weaver (2003); Stewart et al. (2004); Gowrishankar et al. (2013)] and equivalent circuit models [Schoenbach et al. (2004); Ramos et al. (2003)]. The effect of surface charge and membrane conductivity was studied on the induced potential in spherical and non-spherical cell geometries by Kotnik et al., [Kotnik and Miklavčič (2006)] and Mezeme et al., [Mezeme and Brosseau (2010)].

In the last decade, the theoretical approaches of studying the influence of external EF on the dynamics of hMSCs have begun. Although experiments have shown that the external EF affects cellular processes, the theoretical approaches have mainly focused on the collective dynamics of stem cells [Renardy et al. (2018); Farooqi et al. (2019); Sarkar et al. (2019); Paździorek (2014); MacArthur (2014); Lei et al. (2014); Tonge et al. (2010)]. Such approaches consider the stem cell population as a compartment and do not resolve the dependency of processes of individual cells on the external factors. To the best of our knowledge, any of the existing mathematical models have not incorporated the cellular responses of interaction with EF distribution in the cell compartments.

In this context, we investigate the influence of applied EFs on the dynamics of an *in vitro* culture of hMSCs in a TC setup (see Sections 2.2, 2.3). Our mean-field theoretical framework takes into account processes at the scale of an individual stem cell and describes the dynamics of a stem cell population (see Section 3.1 and 3.2). We compare our theory with experimental results reported by Hess et al., and provide a quantitative explanation for the observed behavior of the total number of cells and the total alkaline phosphatase (ALP) activity over time.

## 2 MATERIALS AND METHODS

Our data-driven modeling is based on previous experiments by Hess et al. We use the time dependent experimental data from Hess et al., to study the effect of EFs on hMSC proliferation and differentiation. In the following subsections we recapitulate the experimental TC setup and quantification procedure introduced in Hess et al. We then discuss the corresponding experimental results of the total number of cells and the total ALP activity in the stimulation chamber, which forms the basis for our general theoretical framework.

### 2.1 Electrical stimulation with TC-induced electrical field

The hMSCs were isolated from bone marrow aspirates of 3 healthy male donors between the age of 20 to 40 years old (for more details on isolation and expansion of cells in [Hess et al. (2012b)]). In a spinner flask containing expansion medium (exm, Dulbecco’s modified Eagle’s medium with 10% fetal bovine serum and 100 I.U.*/*mL penicillin-streptomycin), about 50,000 hMSCs were seeded on a collagen-coated polycaprolactone (PCL) disc-shaped scaffold at 37 °C with 7% CO_2_ for 24 h [Hess et al.]. Subsequently, the PCL-scaffolds with hMSCs were transferred to a cultivation chamber of the transformer-like coupling (TC) system previously described [Hess et al.] and prepared for electrical stimulation. In each cultivation chamber, only PLC-scaffolds seeded with the same donor were allowed, so as to avoid side-effects induced by endocrine signaling between hMSC from different donors. Next, 100 ml osteogenic differentiation medium (osm) composed of exm supplemented with 10 nM dexamethasone, 0.2 *µ*M ascorbic acid and 10 mM *β*-glycerophosphate (all Sigma Aldrich); was added to each cultivation chamber and incubated at 37 °C, 7% CO_2_. Further, medium change was performed every 4 days over the entire course of cultivation. An electrical stimulation regime with rectangular pulses (7 ms, 3.6 mV/cm, 10 Hz) was applied intermittently (4 h stimulation, 4 h pause) on the samples over 28 days [Hess et al. (2012b,a)]. To ensure a homogeneous EF for the cell culture, the cells are seeded on the long arms of the chamber where the electrical field was uniform. Our FEM simulation of the chamber confirms the same (see Figure 1(b)). Corresponding negative controls without electrical stimulation were set up in identical cultivation chambers, but were not connected to the transformer core.

**Figure 1.**
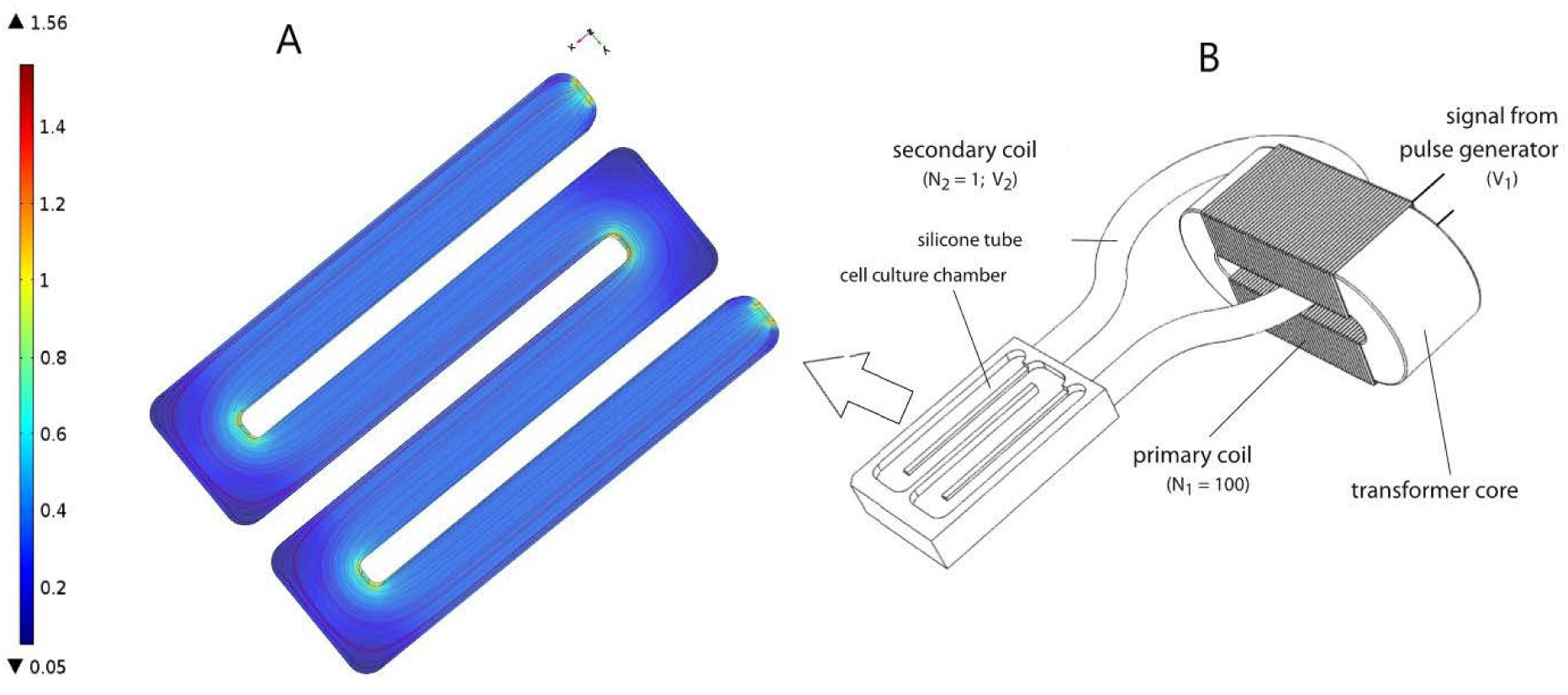
(A) Finite Element Model of cell culture chamber with the electrical field as described in [Hess et al. (2012b)]. FEM model was performed using COMSOL Multiphysics 4.2a®**(B)** Schematic representation of TC, as described in [Hess et al. (2012b)]. *Figure obtained with permission from [Hess et al. (2012b)]*.

### 2.2 Total number of cells and total ALP activity in the TC apparatus

To study the influence of EFs on stem cell dynamics, cell proliferation and cell differentiation were quantified using standard calorimetric measurement protocol. hMSC proliferation and differentiation were determined via lactate dehydrogenase (LDH) and ALP assay, respectively. Experimental data was recorded on 7 d, 14 d, 21 d and 28 d after the electrical stimulation regime was applied. Four samples from each condition (control and electrically stimulated) were collected and stored in −80 °C for analyses later as a whole. Briefly, ALP and LDH assays were performed. To prepare the samples for analysis, they were thawed on ice for 30 min, followed by cell lysis for 50 min in cold lysis buffer consist of 1% w/v Triton X-100 / Phosphate buffer saline (PBS). To determine ALP activity at each time point, 25 *µ*l cell lysate was added to 125 *µ*l ALP substrate consisting 1 mg/ml p-nitorphenyl phosphate (Sigma Aldrich), 0.1 M diethanolamine, 1 mM MgCl_2_ and 0.1% w/v Triton X-100/PBS, pH 9.8. The reaction was prepared in 96-well microplate, incubated at 37 °C for 30 min and stopped with 73 *µ*l NaOH. This is followed by centrifugation at 16 000 g for 10 min and 170 *µ*l of supernatant from individual well was transferred to a new 96-well microplate. The absorbance was measured on TECAN microplate reader at 405 nm and corresponding negative controls had used lysis buffer instead of cell lysate. ALP activity is interpreted as *µ*mol para-nitrophenol (pNP) per 10^6^ cells. To determine the cell number present in each scaffold over time, 50 *µ*l of cell lysate was added to equal volume of LDH substrate (Takara, France) in a 96-well microplate and incubated at room temperature for 5 min. The reaction was stopped by adding 50 *µ*l 0.5 M HCl to each well and the absorbance was measured on TECAN microplate reader at 492 nm. The cell numbers will be determined by correlating the measured values against a calibration curve derived with defined number of hMSCs. For both assays, the measurements were done in triplicates to increase the accuracy.

### 2.3 Experimental Results

Cell proliferation, indicated by the change in the total number of cells over time, shows continuous increase for 28 days, in both the electrically stimulated samples and the non-stimulated control samples, Figure (2A). Statistical analysis showed no detectable differences in the total number of cells between stimulated and non-stimulated samples [Hess et al.]. The total ALP activity in all the hMSCs in the scaffold of the stimulation chamber showed an initial increase, reaching its maximum around 14 days, followed by a decrease over time until 28 days. The statistical analysis showed significant difference between the electrically stimulated and non-stimulated control samples. The ALP activity in the electrically stimulated samples was 30 % higher than the non-stimulated control samples. This indicates a role of the applied EFs in the differentiation process.

**Figure 2.**
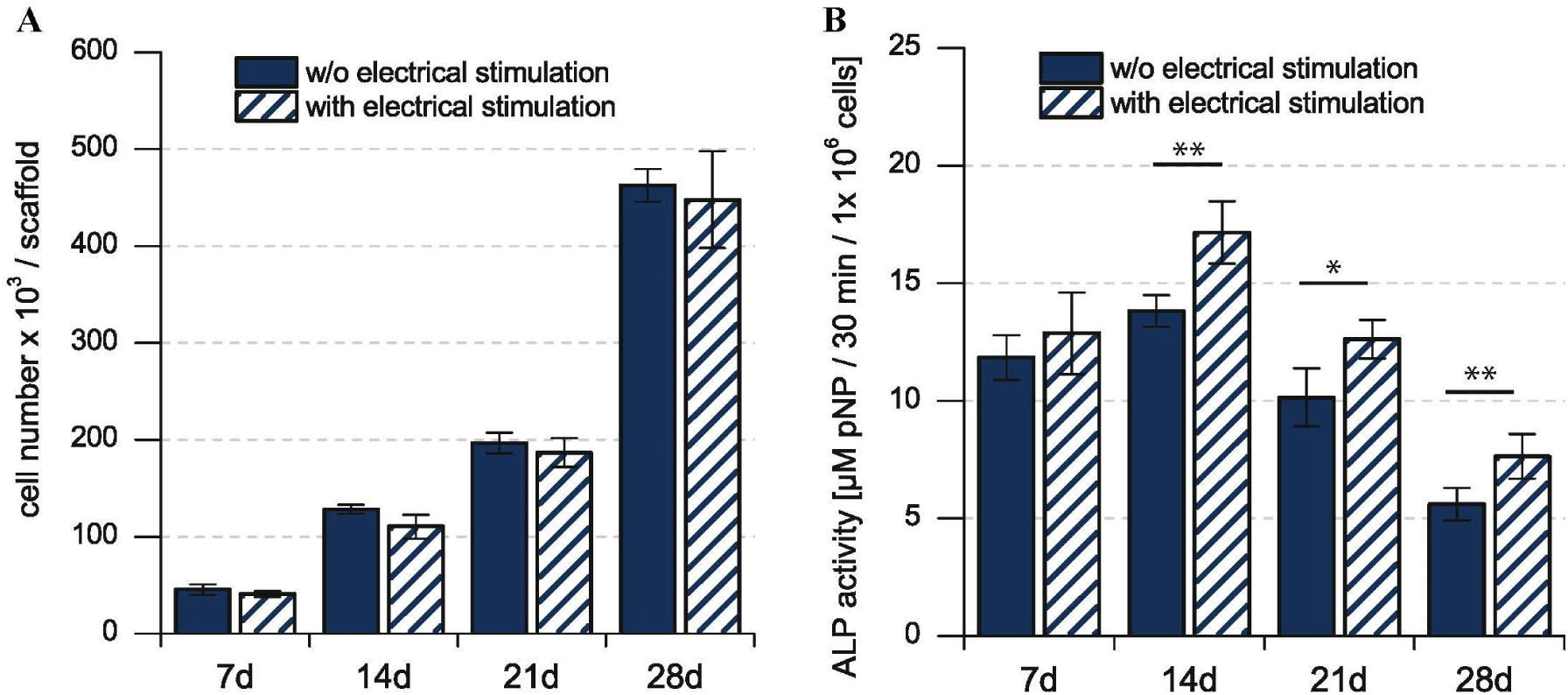
(A) Total number of mesenchymal stem cells in the cell culture. **(B)** Biochemical analysis of the total ALP activity in the cell culture. Total number of number of hMSCs is indicative of the stem cell proliferation, whereas the total ALP activity is indicative of the stem cell differentiation. *\** is used for p *<* 0.05, *\*\** for p *<* 0.01. *Figure obtained with permission from [Hess et al.]*

## 3 MATHEMATICAL MODELING

Based on the experimental data, discussed in the previous section, we make the following observations. First, the non-stimulated control samples show different time dependent behaviors for the total number of cells and the total ALP activity. Second, the applied EF significantly influences only the time dependent behavior of the total ALP activity. In order to provide a quantitative explanation for these observations, we formulated a general theoretical description of stem cell dynamics.

### 3.1 General Theoretical Framework for Stem Cell Dynamics

In our mean-field model, the time-dependent behavior of the stem cell population augments from the processes at the scale of a single stem cell. An individual stem cell undergoes division, giving rise to new cells and thus sustaining the stem cell population. The ALP activity of a stem cell is maintained by the intracellular biochemical processes. Subsequently, a stem cell leaves the stem cell population due to terminal differentiation. Taking these processes into account at the scale of an individual cell, we describe the state of the mesenchymal stem cell population by *n*(*a, t*), where *a* is the ALP activity of a cell in the stem cell population at time *t*. Precisely, *n*(*a, t*)Δ*a* is the total number of cells with ALP activity in the range *a* and *a* + Δ*a* at the time *t*. Generally, *n* can depend on multiple variables besides intracellular ALP activity, such as the cell size, the ALP gene expression of the cell, the EF strength experienced by the cell, the orientation of the cell with respect to the applied EF etc. The change in *n* over time reflects the dynamics of individual stem cells.

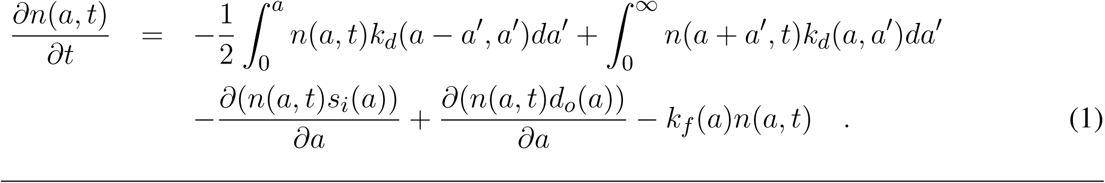

Equation (1) is based on the Smoluchowski equation describing coagulation phenomena [Smoluchowski (1916)]. Similar equations have been studied in a variety of other problems [Foret et al. (2012); Baskaran and Marchetti (2008)]. The first two terms on the right hand side of Equation (1) represent cell divisions in the stem cell population. These cell divisions result in the stem cell proliferation. Such cell divisions occur at the rate *k*_*d*_(*a, a′*), and replace a cell having ALP activity *a* + *a′*, with two daughter cells having ALP activities *a* and *a′*, respectively. These cell divisions described in Equation (1) conserve the ALP activity. In the case of non-conserved cell divisions, both the daughter cells have the same measure of ALP activity as the dividing parent cell. Such cell divisions will be described by only one term, namely, 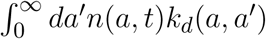 instead of two terms as in Equation (1). The third and the fourth term on the right hand side of Equation (1) represent the flux of ALP activity in the cell due to the intracellular biosynthesis and degradation of ALP, respectively. In our theoretical description we assume the ALP activity of a stem cell to be regulated by the intracellular ALP biosynthesis and degradation processes. For a cell with ALP activity *a, s_i_*(*a*) is the average ALP activity gained per unit time due to ALP biosynthesis and *d*_*o*_(*a*) is the average ALP activity lost per unit time due to ALP degradation. The last term on the right hand side of Equation (1) represents the loss of a cell with ALP activity *a* from the population. Such losses occur at the rate *k*_*f*_ (*a*) due to instantaneous differentiation of a hMSC into a fully differentiated osteoblast cell. Our theoretical framework describes the dynamics of a population of undifferentiated hMSCs, and does not include osteoblasts, i.e. terminally differentiated hMSCs. In our description, cells with measurable ALP activity are classified as undifferentiated mesenchymal stem cells. We assume the osteoblasts to have lower ALP activity, compared to the undifferentiated mesenchymal stem cells. We also assume that the intracellular ALP activity reaches its maximum in the mesenchymal stem cells undergoing differentiation. Now, we introduce two quantities *N* (*t*) and Φ(*t*) as follows,

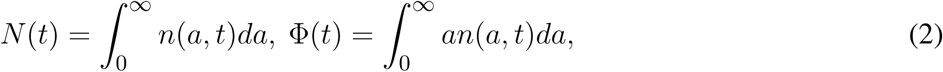

where, *N* (*t*) represents the total number of cells, and Φ(*t*) represents the total ALP activity in the hMSC population at time *t*. The time dependent behaviour of *N* and Φ describe the dynamics of hMSC population as whole. Using Equation (2) and Equation (1), we can write down the balance relations for *N* and Φ, in the case of cell divisions that conserve ALP activity, as follows,

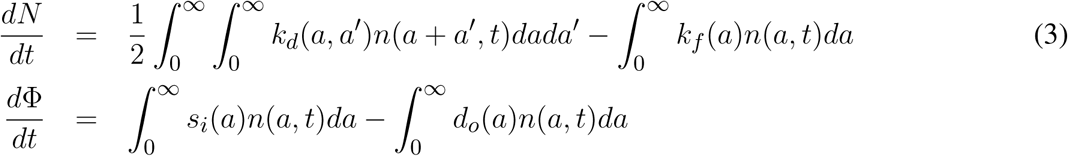

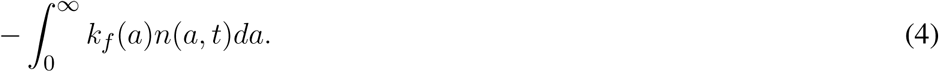

From Equations (3) and (4) we see that the macroscopic quantities *N* and Φ result from the dynamics of individual cells, such as cell division, ALP biosynthesis and degradation governing cellular ALP activity and cell differentiation.

## 4 RESULTS

The experimentally observed time dependent behaviour of *N* and Φ can be explained by a model that includes stem cell proliferation due to cell divisions and osteogenic differentiation. Mesenchymal stem cell differentiation into an osteoblast could either occur instantaneously or proceed gradually giving rise to intermediate pre-osteoblasts with detectable ALP activity. Our theoretical framework distinguishes between these two subtly different processes, which will be discussed in the following.

### 4.1 Progressive Stem Cell Differentiation Model

In this model, stem cell proliferation occurs due to symmetric non-conserved cell divisions. The osteogenic differentiation in this model is due to a gradual decrease in ALP activity via intracellular ALP degradation. The parameter choice for this model is given in Table (1). The cell division in this model occurs at the rate of *k*_*d*_ and results in two daughter cells with the same magnitude of the ALP activity as the parent cell. The stem cell differentiation in this model occurs gradually at the rate of *d*_*o*_, i.e. ALP out-flux due to intracellular ALP degradation is proportional to the cell’s ALP activity, Figure (4). The dynamic equation for *n*(*a, t*) using the parameter choice listed in Table (1) is given by,

**Figure 3.**
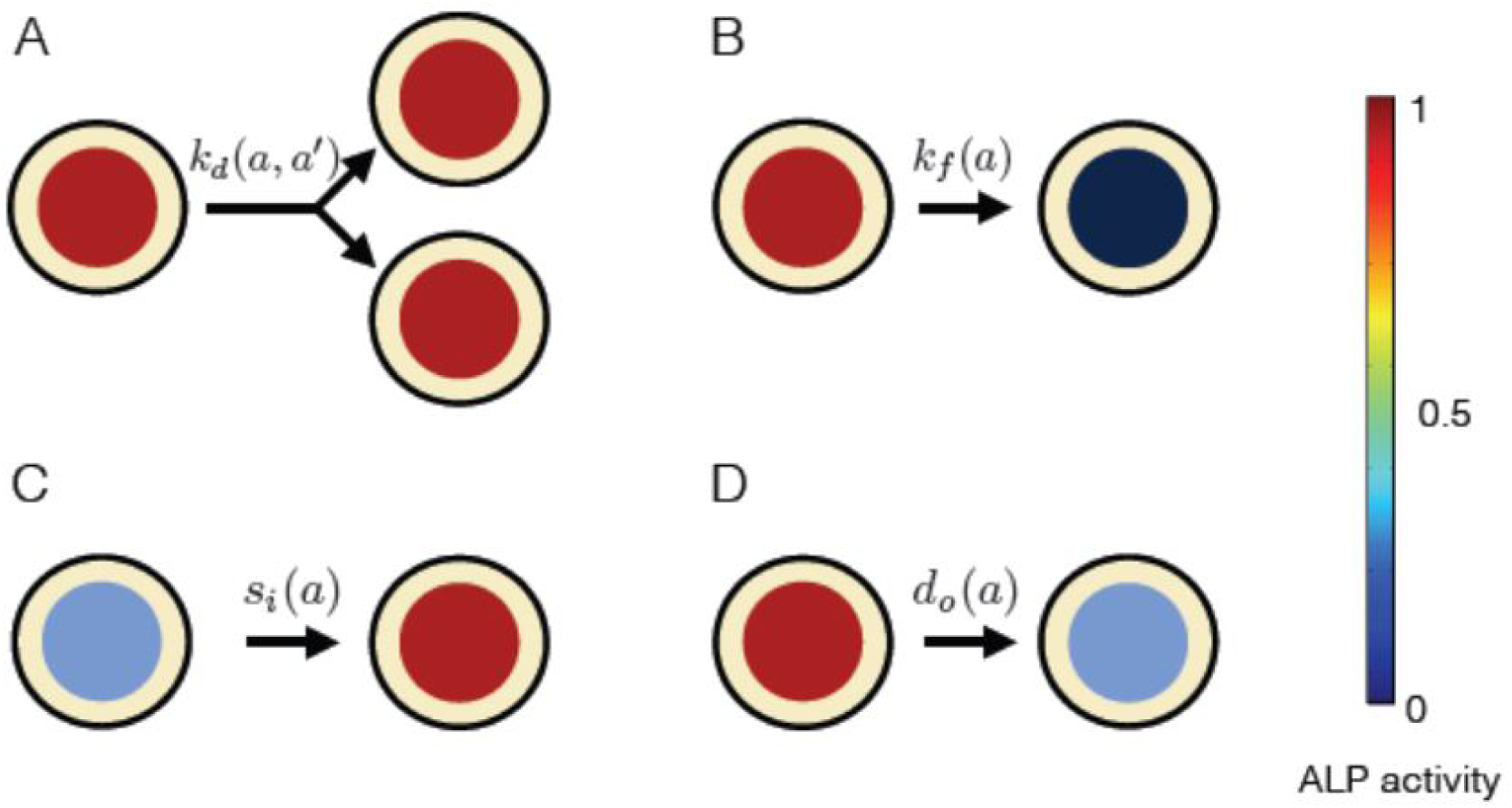
Processes affecting the distribution of ALP activity *n*(*a, t*) in the population of hMSCs. **(A)** When a cell with ALP activity *a* + *a′* divides, it is replaced by two new cells with ALP activity *a* and *a′*. The colored circles represent the level of the ALP activity of the cell, according to the color scale. High level of the ALP activity is represented by red color, whereas no ALP activity is represented by black color.. Such divisions occur at the rate *k*_*d*_(*a, a′*) and result in the stem cell proliferation. **(B)** A cell can instantaneously lose detectable ALP activity because of terminal osteogenic differentiation into an osteoblast. Such processes occur at the rate *k*_*f*_ (*a*). **(C)** ALP activity in a cell can increase due to intracellular ALP synthesis, denoted by influx *s*_*i*_(*a*). **(D)** ALP activity in a cell can decrease due to intracellular ALP degradation, denoted by out-flux *d*_*o*_(*a*).

**Figure 4.**
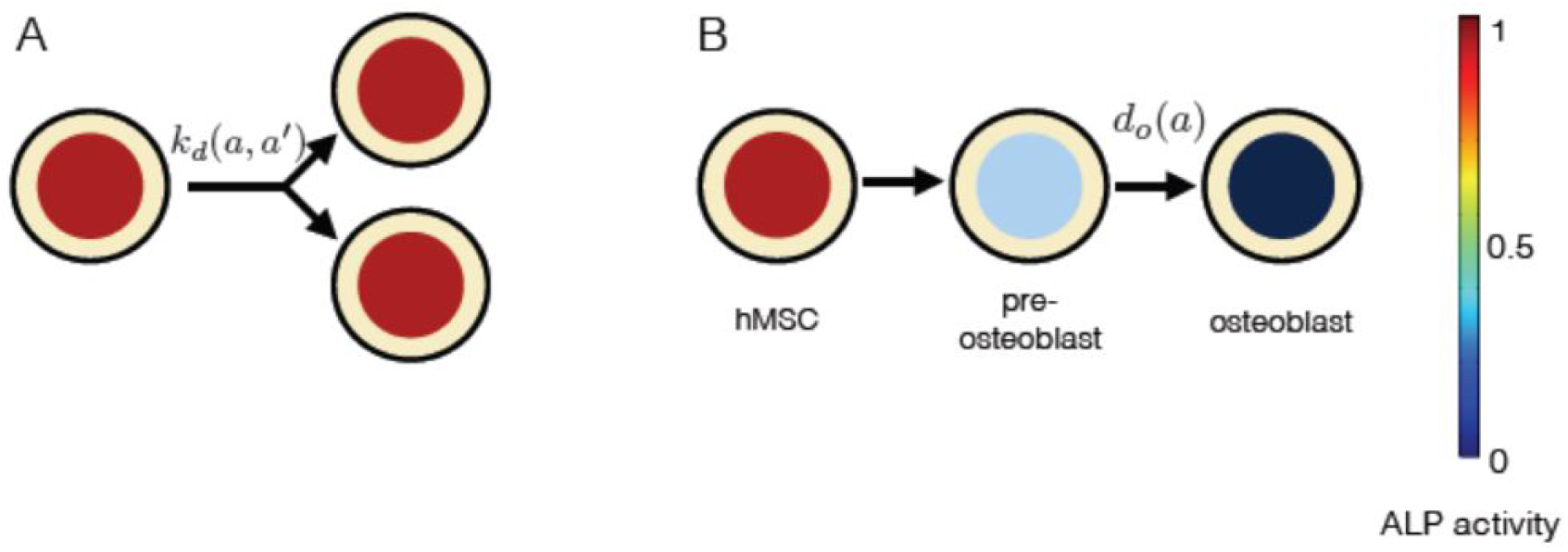
The constituents of progressive stem cell differentiation model are **(A)** Symmetric non-conserved cell division. The division of a stem cell with ALP activity *a* results in two daughter stem cells, each with ALP activity *a*. **(B)** Osteogenic differentiation. The osteogenic differentiation in Model 1 proceeds via gradual loss of ALP activity. Such a process gives rise to temporary states of intermediate pre-osteoblasts.

**Table 1.**
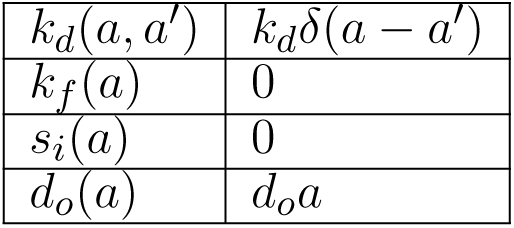
Choice of parameters for progressive stem cell differentiation model. This model includes cell proliferation due to symmetric non-conserved cell divisions and gradual differentiation of a hMSC into an osteoblast cell due to out-flux of ALP activity.

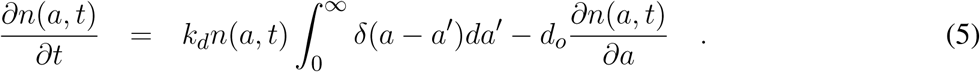

Equation (5) is solved by using the Laplace transformation technique (Supplemental Information) to obtain the balance relations for *N* and Φ. The time rate of change of *N* is,

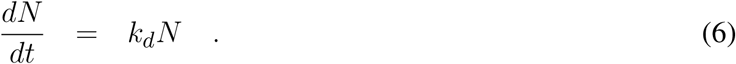

To fit Equation (6) to the experimental data of the total number of stem cells at each time point in the stimulation chamber, we used *k*_*d*_ = 2*/t*. The solution of Equation (6) with this choice of *k*_*d*_ is given by,

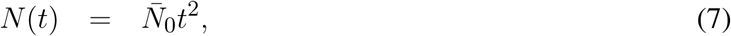

where 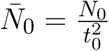 and *N*_0_ is the total number of cells showing ALP activity in the hMSC population at the initial time *t*_0_. The fitting was performed using the least squares fitting method, giving an estimate for 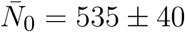, Figure (5A). The dynamic equation for Φ, obtained from Equation (5), is

**Figure 5.**
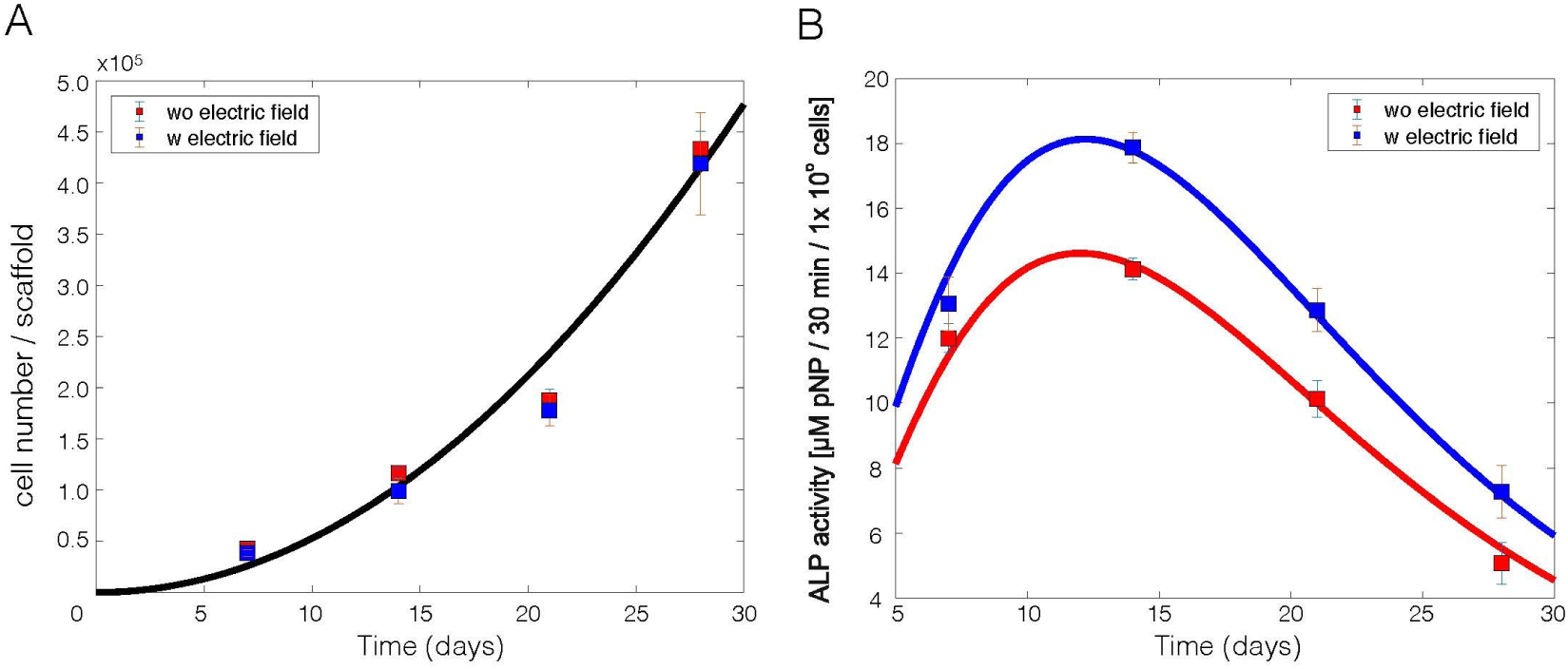
Experimental results for electrically stimulated (blue box) and non-stimulated (red box) cell culture samples after 7, 14, 21 and 28 days. **(A)** The total number of cells in the stimulation chamber. The solid curve is the fit of the analytical solution of Model 1, given by Equation (7), to the experimental data. **(B)** The total ALP activity of the cells in the stimulation chamber. The solid curve is the fit of the analytical solution of progressive stem cell differentiation (PSCD) model, given by Equation (9), to the experimental data. Error-bars show the standard deviation in the experimental data.

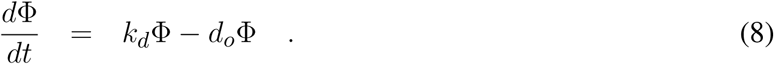

The solution of Equation (8), with our choice of *k*_*d*_, is

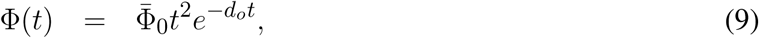

where 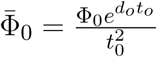 and Φ_0_ is the total ALP activity in all the cells in the hMSC population at the initial time *t*_0_. Equation (9) was fit to the experimental data of the total ALP activity at each time point in the stimulation chamber, Figure (5B). Since the statistical analysis showed a significant difference between the non-stimulated control and electrically stimulated samples, Figure (2(B)), we performed data fitting of each sample separately, Figure (5(B)). The parameter values of the function, given by Equation (9), obtained as a result of the fit to the experimental data are given in Table (2).

**Table 2.**
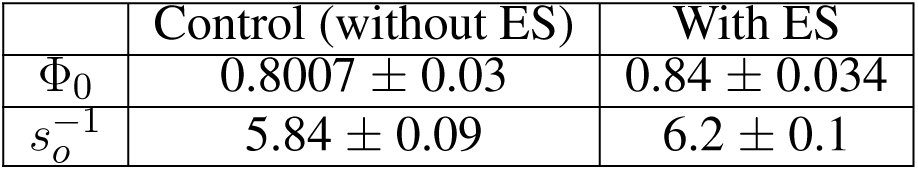
Parameter values obtained as a result of the fit of Equation (9) to the experimental total ALP activity of stem cells in the stimulation chamber.

|  | Control (without ES) | With ES |
| --- | --- | --- |
| $\Phi_0$ | $0.8007 \pm 0.03$ | $0.84 \pm 0.034$ |
| $s_o^{-1}$ | $5.84 \pm 0.09$ | $6.2 \pm 0.1$ |

### 4.2 Instantaneous Stem Cell Differentiation Model

Instantaneous stem cell differentiation model (ISCD) includes stem cell proliferation due to symmetric non-conserved cell divisions, similar to PSCD model. However, the difference between the two models lies in the precise mechanism of osteogenic differentiation. In ISCD model, the differentiation of a stem cell into an osteoblast cell occurs instantaneously, resulting in the total loss of ALP activity in the differentiated osteoblast cell, Figure (6). In this model, such a sudden loss of a cell with ALP activity might also imply apoptosis. The parameter choice for ISCD model is given in Table (3). The dynamic equation for *n*(*a, t*) using the parameter choice listed in Table (3) is given by,

**Table 3.**
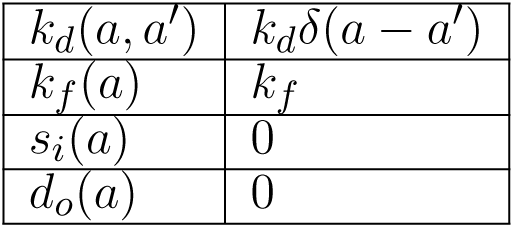
Choice of parameters for instantaneous stem cell differentiation model. This model includes cell proliferation due to symmetric non-conserved cell divisions and instantaneous differentiation of a hMSC into osteoblast cell.

**Figure 6.**
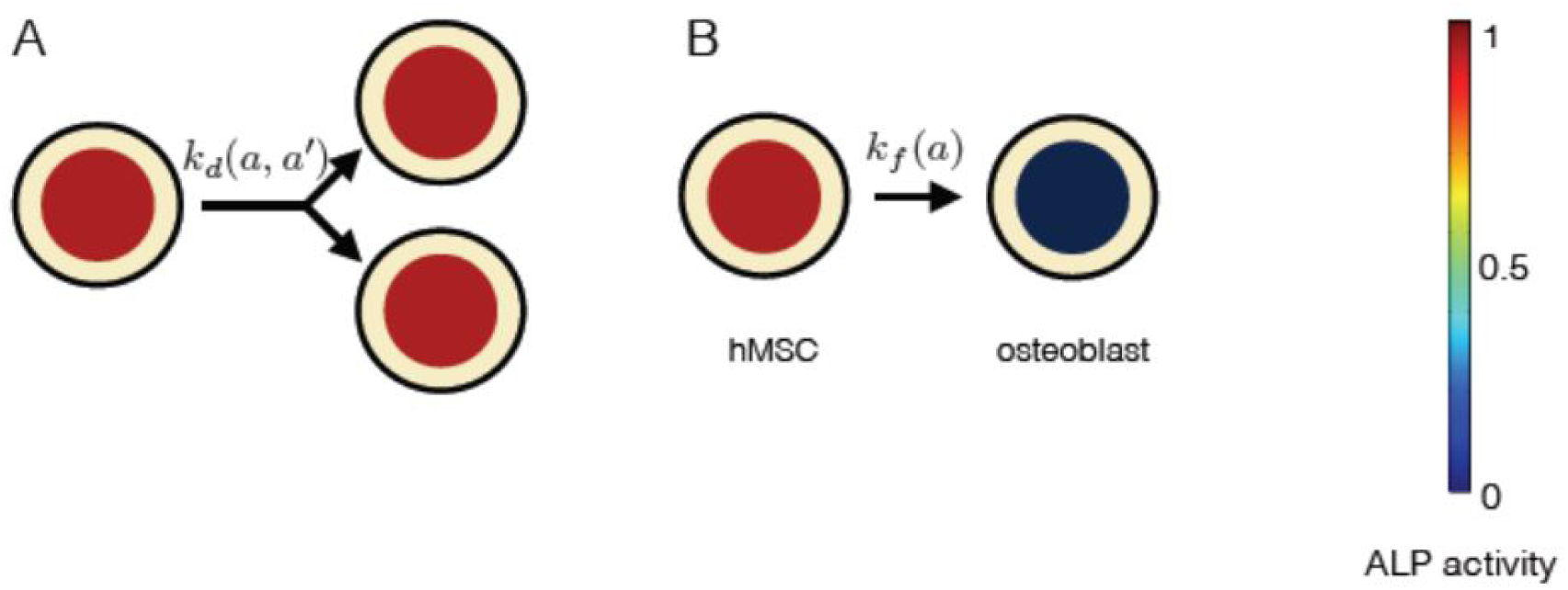
The constituents of progressive stem cell differentiation model are **(A)** Symmetric non-conserved cell division. The division of a stem cell with ALP activity *a* results in two daughter stem cells, each with ALP activity *a*. **(B)**The osteogenic differentiation in instantaneous stem cell differentiation model occurs instantaneously via sudden loss of a cell’s ALP activity. Such a process instantaneously gives rise to an osteoblast cell with no ALP activity.

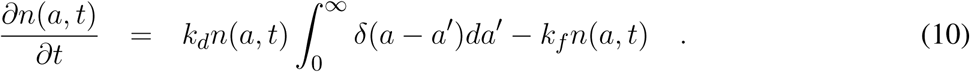

The change of *N* and Φ over time is,

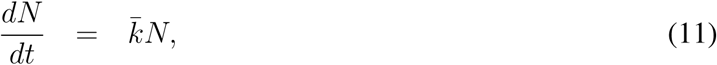

and,

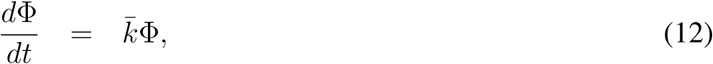

respectively, where 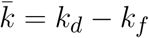.

Since Equation (11) and Equation (12) have exactly the same form, their solutions also have exactly the same functional form. The solution of Equation (11) and Equation (12), by choosing 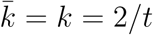 as in Model 1, we get,

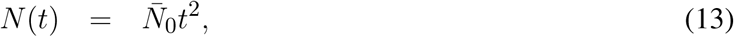

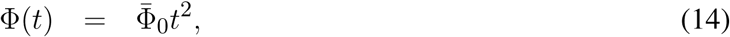

where, 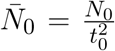 and 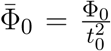. *N*_0_ and Φ_0_ are as defined in Model 1. The ISCD model describes the experimental data for the total number of cells, but it fails to capture the non-monotonic time dependent behaviour of the experimental data for the total ALP activity. Equation (14) shows a continuous increase of the total ALP activity for all time points, whereas experimental data, for both the control and stimulated samples, show an increase in the total ALP activity only up to 14 days, Figure (2B). After 14 days, the total ALP activity shows a continuous decrease till 28 days in both control and stimulated samples which is not captured by this model, Figure (2B).

## 5 DISCUSSION

In this study we developed a general theoretical framework to describe how applied EFs influence the stem cell dynamics. Our mean-field description of stem cell dynamics augments from elementary processes such as stem cell division, differentiation and intracellular regulation of ALP activity. Current theoretical approaches to the study of stem cell dynamics are based on biochemical assays that consider stem cell population as a whole and do not resolve processes at the scale of individual cells. Such approaches ignore the discrete nature of the stem cell population consisting of many individual cells. Our theoretical framework takes into account processes governing the dynamics of individual cells in the stem cell population. The advantage of our general theory is that it allows for studying the influence of various factors on the rates of cellular processes. In addition, our theoretical framework can distinguish between different mechanisms through which cellular processes occur. We tested our theory with *in vitro* electrical stimulation experiments by Hess et al. [Hess et al. (2012b)]. We show that our first model, derived from our general theory, PSCD model, can fully describe the time dependent behaviours of the total number of ALP expressing hMSCs and the total ALP activity in the stimulation chamber. In this model, stem cell proliferation is due to symmetric non-conserved cell divisions and stem cell differentiation occurs via gradual loss of the ALP activity in the stem cells. In the second model, referred to as the instantaneous stem cell differentiation model, we studied the cell differentiation due to the sudden loss of ALP activity, and its effect on the stem cell dynamics. The rates at which the cell division and the cell differentiation occur, in these models, depend on the ALP activity of the stem cell.

Our analysis reveals a negative correlation between the stem cell proliferation rate and the cell number density, Figure (7(A)). The coupling between the cell proliferation rate and the cell density could either be due to density-dependent inter-cellular signaling or mechanical compression, or both [Eyckmans et al. (2012); Najafabadi et al. (2016)]. Recent experiments have shown that the *in vitro* osteogenic differentiation is associated with the processing of type-1 collagen and progressive deposition of the extracellular collagen matrix [Hanna et al. (2018)]. The deposition of the extracellular matrix over time might restrict cells from growing and dividing. This could explain the dependence of cell proliferation rate on the cell density, as our analysis suggests. The density-dependent cell division rate has been explored in other context of cellular systems as well [Recho et al. (2016); Hoffmann et al. (2011)].

**Figure 7.**
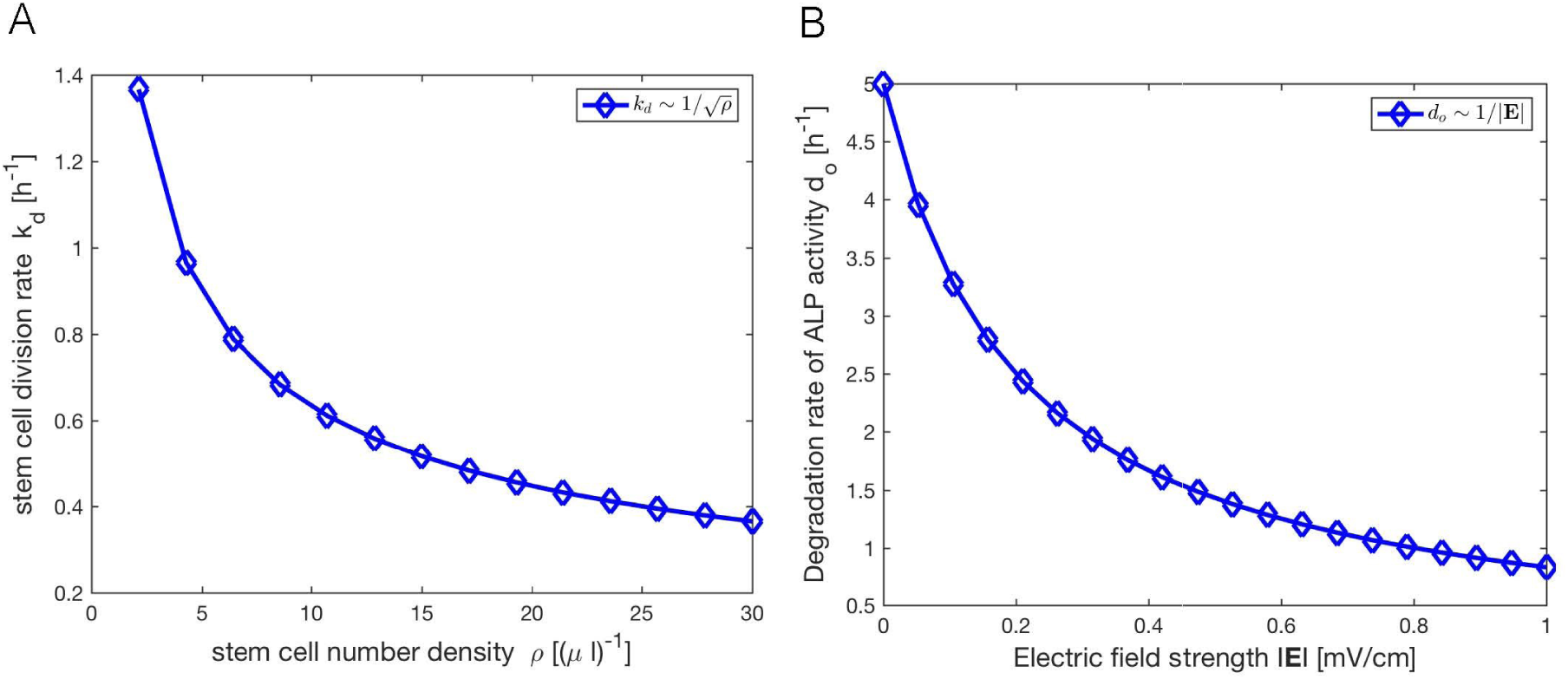
The dynamics of hMSCs is influenced by the cell number density and the applied EF. **(A)** The analysis presented in this study reveals a negative correlation between the divison rate *k*_*d*_ of the stem cells and the stem cell number density *ρ*. The stem cell number density is given by the relation *ρ* = *N/V*, where *N* is the total number of stem cells at the time *t* in the scaffold of the stimuation chamber, and *V* is the volume of scaffold, which is fixed. **(B)** The analysis also suggests that the degradation rate of the ALP activity *d*_*o*_ of a cell is inversely proportional to the strength of the applied EF |**E**|. *d*_*o*_ indicates the rate of osteogenic differentiation.

Our results show that the applied EF influences stem cell differentiation rather than stem cell proliferation, which confirms the experimental result of Hess et al. [Hess et al. (2012b)]. We found that the rate of degradation of the ALP activity is inversely proportional to the applied EF strength, Figure (7(B)). In order to precisely quantify the dependency of the stem cell differentiation on the applied EF, further studies of stimulation of hMSCs with varying field strengths are needed. A comparison of our two precisive models with the experiments suggests that the stem cell differentiation occurs gradually, as described in the progressive stem cell differentiation model. This mechanism of osteogenic differentiation gives rise to pre-osteoblast cells, which confirms the experimental results of Rutkovskiy et al. (2016)].

## 6 CONCLUSION

We draw the following conclusions from our analysis presented in this study. First, despite the complexity of the process, reflected in the multiplicity in its regulatory steps, we show that the stem cell dynamics can be understood by a simple description that captures vital processes. Secondly, our analysis shows that the applied EFs predominantly influence stem cell differentiation. Thirdly, we show that the progressive stem cell differentiation model thoroughly describes the experimental results of Hess et al. [Hess et al. (2012b)]. This model suggests that the osteogenic differentiation of hMSCs progresses, gradually giving rise to pre-osteoblast cells. Fourthly, our analysis shows that the stem cell division rate is cell density-dependant. Finally, our framework allows to measure the rates of cellular processes and estimates their dependency on external EF. This method is robust and sensitive since the time dependent macroscopic quantities result from an average over many individual cells and multiple experimental repetitions. This framework could serve as a tool to study the influence of external factors on stem cell dynamics through genetic and chemical perturbation of various cellular processes.

## 7 PERMISSION TO REUSE AND COPYRIGHT

Figures 1B, 3A and 3B are reused with permission from [Hess et al. (2012b)].

## AUTHOR CONTRIBUTIONS

RA and UvR have conceptualized the study; RA has designed the project, supervised the work including data analysis and model development; JD developed the mean-field model and derived analytical results; JD and RA worked on the simple model for dynamics MSCs; JD performed data fitting; RA has performed the FEM simulations; JD, PL, UvR and RA wrote sections of the manuscript. RA critically revised the manuscript and took the responsibility for the integrity of the study as a whole. All authors contributed to manuscript revision, read and approved the submitted version.

## ACKNOWLEDGMENTS

We are grateful to Dr. Ricarda Hess and Prof. Dieter Scharnweber for providing the experimental data for this mathematical modeling. We thank Prof. Hans-Peter Wiesmann for permitting PL to work on our analysis and acknowledge the financial support from the DFG Transregio 67 (Project A3). We acknowledge Dr. Kiran Sriperumbudur for his help in revising the paper.

## SUPPLEMENTAL DATA

The supplementary material for this article can be found at:

## REFERENCES

(2006). Matrix Elasticity Directs Stem Cell Lineage Specification. Cell 126, 677–689. doi: 10.1016/j.cell.2006.06.044

(2009). Novel Effect of Biphasic Electric Current on In Vitro Osteogenesis and Cytokine Production in Human Mesenchymal Stromal Cells. Tissue Engineering Part A 15, 2411–2422. doi: 10.1089/ten.tea.2008.0554

Banks, T. A., Luckman, P. S. B., Frith, J. E., and Cooper-White, J. J. (2015). Effects of electric fields on human mesenchymal stem cell behaviour and morphology using a novel multichannel device. Integrative Biology 7, 693–712. doi: 10.1039/c4ib00297k

Baskaran, A. and Marchetti, M. C. (2008). Enhanced diffusion and ordering of self-propelled rods. Physical Review Letters, 268101 doi: 10.1103/PhysRevLett.101.268101

Chang, W. H.-S., Chen, L.-T., Sun, J.-S., and Lin, F.-H. (2004). Effect of pulse-burst electromagnetic field stimulation on osteoblast cell activities. Bioelectromagnetics 25, 457–465. doi: 10.1002/bem.20016

Ciombor, D. M. and Aaron, R. K. (1993). Influence of electromagnetic fields on endochondral bone formation. Journal of Cellular Biochemistry 52, 37–41. doi: 10.1002/jcb.240520106

De-Leon, S. B.-T. and Davidson, E. H. (2007). Gene Regulation: Gene Control Network in Development. Annual Review of Biophysics and Biomolecular Structure 36, 191–212. doi: 10.1146/annurev.biophys.35.040405.102002

Dingal, P. C. D. P., Wells, R. G., and Discher, D. E. (2014). Simple insoluble cues specify stem cell differentiation. Proceedings of the National Academy of Sciences 111, 18104–18105. doi: 10.1073/pnas.1421562112

Eyckmans, J., Lin, G. L., and Chen, C. S. (2012). Adhesive and mechanical regulation of mesenchymal stem cell differentiation in human bone marrow and periosteum-derived progenitor cells. Biology Open 1, 1058–1068. doi: 10.1242/bio.20122162

Farooqi, A. R., Bader, R., and Van Rienen, U. (2019). Numerical Study on Electromechanics in Cartilage Tissue with Respect to Its Electrical Properties, 152–166 doi: 10.1089/ten.teb.2018.0214

Foret, L., Dawson, J. E., Villaseñor, R., Collinet, C., Deutsch, A., Brusch, L., et al. (2012). A general theoretical framework to infer endosomal network dynamics from quantitative image analysis. Current Biology, 1381–90 doi: 10.1016/j.cub.2012.06.021

Fricke, H. (1953). The Electric Permittivity of a Dilute Suspension of Membrane-Covered Ellipsoids. Journal of Applied Physics 24, 644–646. doi: 10.1063/1.1721343

Funk, R. H. W. (2015). Endogenous electric fields as guiding cue for cell migration. Frontiers in Physiology 6, 143. doi: 10.3389/fphys.2015.00143

Gimsa, J. and Wachner, D. (2001a). Analytical description of the transmembrane voltage induced on arbitrarily oriented ellipsoidal and cylindrical cells. Biophysical Journal 81, 1888–1896. doi: 10.1016/S0006-3495(01)75840-7

Gimsa, J. and Wachner, D. (2001b). On the analytical description of transmembrane voltage induced on spheroidal cells with zero membrane conductance. European Biophysics Journal 30, 463–466. doi: 10.1007/s002490100162

Gowrishankar, T. R., Smith, K. C., and Weaver, J. C. (2013). Transport-based biophysical system models of cells for quantitatively describing responses to electric fields 101, 505–517. doi: 10.1109/JPROC.2012.2200289

Gowrishankar, T. R. and Weaver, J. C. (2003). An approach to electrical modeling of single and multiple cells. Proceedings of the National Academy of Sciences 100, 3203–3208. doi: 10.1073/pnas.0636434100

Grosse, C. and Schwan, H. P. (1992). Cellular membrane potentials induced by alternating fields. Biophysical Journal 63, 1632–1642. doi: 10.1016/S0006-3495(92)81740-X

Hanna, H., Mir, L. M., and Andre, F. M. (2018). In vitro osteoblastic differentiation of mesenchymal stem cells generates cell layers with distinct properties. Stem Cell Research and Therapy doi: 10.1186/s13287-018-0942-x

Hartig, M., Joos, U., and Wiesmann, H.-P. (2000). Capacitively coupled electric fields accelerate proliferation of osteoblast-like primary cells and increase bone extracellular matrix formation in vitro. European Biophysics Journal 29, 499–506. doi: 10.1007/s002490000100

Hess, R., Jaeschke, A., Neubert, H., Hintze, V., Moeller, S., Schnabelrauch, M., et al. (2012a). Synergistic effect of defined artificial extracellular matrices and pulsed electric fields on osteogenic differentiation of human MSCs. Biomaterials 33, 8975–8985. doi: 10.1016/j.biomaterials.2012.08.056

Hess, R., Neubert, H., Seifert, A., Bierbaum, S., Hart, D. A., and Scharnweber, D. (2012b). A Novel Approach for In Vitro Studies Applying Electrical Fields to Cell Cultures by Transformer-Like Coupling. Cell Biochemistry and Biophysics doi: 10.1007/s12013-012-9388-4

Hoffmann, M., Kuska, J. P., Zscharnack, M., Loeffler, M., and Galle, J. (2011). Spatial organization of mesenchymal stem cells in vitro-results from a new individual cell-based model with podia. PLoS ONE doi: 10.1371/journal.pone.0021960

Jansen, J. H., van der Jagt, O. P., Punt, B. J., Verhaar, J. A., van Leeuwen, J. P., Weinans, H., et al. (2010). Stimulation of osteogenic differentiation in human osteoprogenitor cells by pulsed electromagnetic fields: an in vitro study. BMC Musculoskeletal Disorders 11, 188. doi: 10.1186/1471-2474-11-188

Kotnik, T., Bobanović, F., and Miklavčič, D. (1997). Sensitivity of transmembrane voltage induced by applied electric fields - A theoretical analysis. Bioelectrochemistry and Bioenergetics 43, 285–291. doi: 10.1016/S0302-4598(97)00023-8

Kotnik, T. and Miklavčič, D. (2006). Theoretical evaluation of voltage inducement on internal membranes of biological cells exposed to electric fields. Biophysical Journal 90, 480–491. doi: 10.1529/biophysj.105.070771

Lei, J., Levin, S. A., and Nie, Q. (2014). Mathematical model of adult stem cell regeneration with cross-talk between genetic and epigenetic regulation. Proceedings of the National Academy of Sciences of the United States of America, E880–E887 doi: 10.1073/pnas.1324267111

Lin, H.-Y. and Lin, Y.-J. (2011). In vitro effects of low frequency electromagnetic fields on osteoblast proliferation and maturation in an inflammatory environment. Bioelectromagnetics 32, 552–560. doi: 10.1002/bem.20668

Lohmann, C. H., Schwartz, Z., Liu, Y., Guerkov, H., Dean, D. D., Simon, B., et al. (2000). Pulsed electromagnetic field stimulation of MG63 osteoblast-like cells affects differentiation and local factor production. Journal of Orthopaedic Research 18, 637–646. doi: 10.1002/jor.1100180417

MacArthur, B. D. (2014). Collective dynamics of stem cell populations, 3653–3654 doi: 10.1073/pnas.1401030111

Maswiwat, K., Wachner, D., and Gimsa, J. (2008). Effects of cell orientation and electric field frequency on the transmembrane potential induced in ellipsoidal cells. Bioelectrochemistry 74, 130–141. doi: 10.1016/j.bioelechem.2008.06.001

Meny, I., Burais, N., Buret, F., and Nicolas, L. (2007). Finite-element modeling of cell exposed to harmonic and transient electric fields. In IEEE Transactions on Magnetics. vol. 43, 1773–1776. doi: 10.1109/TMAG.2007.892517

Mezeme, M. E. and Brosseau, C. (2010). Time-varying electric field induced transmembrane potential of a core-shell model of biological cells. Journal of Applied Physics 108. doi: 10.1063/1.3456163

Miller, C. and Henriquez, C. (1988). Three-dimensional finite element solution for biopotentials: erythrocyte in an applied field. IEEE Transactions on Biomedical Engineering 35, 712–718

Miyamoto, H., Sawaji, Y., Iwaki, T., Masaoka, T., Fukada, E., Date, M., et al. (2019). Intermittent pulsed electromagnetic field stimulation activates the mTOR pathway and stimulates the proliferation of osteoblast-like cells. Bioelectromagnetics 40, 412–421. doi: 10.1002/bem.22207

Mousavi, S. J. and Hamdy Doweidar, M. (2015). Role of Mechanical Cues in Cell Differentiation and Proliferation: A 3D Numerical Model. PLoS One 10, e0124529. doi: 10.1371/journal.pone.0124529

Najafabadi, M. M., Bayati, V., Orazizadeh, M., Hashemitabar, M., and Absalan, F. (2016). Impact of Cell Density on Differentiation Efficiency of Rat Adipose-derived Stem Cells into Schwann-like Cells. International Journal of Stem Cells 9, 213–220. doi: 10.15283/ijsc16031

Paździorek, P. R. (2014). Mathematical Model of Stem Cell Differentiation and Tissue Regeneration with Stochastic Noise. Bulletin of Mathematical Biology, 1642–1669 doi: 10.1007/s11538-014-9971-5

Petecchia, L., Sbrana, F., Utzeri, R., Vercellino, M., Usai, C., Visai, L., et al. (2015). Electro-magnetic field promotes osteogenic differentiation of BM-hMSCs through a selective action on Ca2+-related mechanisms. Scientific Reports 5, 13856. doi: 10.1038/srep13856

Rajnicek, A. M., Foubister, L. E., and McCaig, C. D. (2008). Alignment of corneal and lens epithelial cells by co-operative effects of substratum topography and DC electric fields. Biomaterials 29, 2082–2095. doi: 10.1016/j.biomaterials.2008.01.015

Ralston, A. (2008). Gene Expression Regulates Cell Differentiation. Nature education

Ramos, A., Raizer, A., and Marques, J. L. (2003). A new computational approach for electrical analysis of biological tissues. Bioelectrochemistry 59, 73–84. doi: 10.1016/S1567-5394(03)00004-5

Recho, P., Ranft, J., and Marcq, P. (2016). One-dimensional collective migration of a proliferating cell monolayer. Soft Matter, 2381–91 doi: 10.1039/c5sm02857d

Renardy, M., Jilkine, A., Shahriyari, L., and Chou, C. S. (2018). Control of cell fraction and population recovery during tissue regeneration in stem cell lineages. Journal of Theoretical Biology, 33–50 doi: 10.1016/j.jtbi.2018.02.017

Rohde, M., Ziebart, J., Kirschstein, T., Sellmann, T., Porath, K., Kühl, F., et al. (2019). Human Osteoblast Migration in DC Electrical Fields Depends on Store Operated Ca2+-Release and Is Correlated to Upregulation of Stretch-Activated TRPM7 Channels. Frontiers in Bioengineering and Biotechnology 7, 422. doi: 10.3389/fbioe.2019.00422

Rutkovskiy, A., Stensløkken, K.-O., and Vaage, I. J. (2016). Osteoblast Differentiation at a Glance. Medical Science Monitor Basic Research, 95–106 doi: 10.12659/msmbr.901142

Sarkar, N., Prost, J., and Jülicher, F. (2019). Field induced cell proliferation and death in a model epithelium. New Journal of Physics doi: 10.1088/1367-2630/ab0a8d

Schemitsch, E. and Kuzyk, P. (2009). The science of electrical stimulation therapy for fracture healing. Indian Journal of Orthopaedics 43, 127. doi: 10.4103/0019-5413.50846

Schoenbach, K. H., Joshi, R. P., Kolb, J. F., Chen, N., Stacey, M., Blackmore, P. F., et al. (2004). Ultrashort electrical pulses open a new gateway into biological cells. In Proceedings of the IEEE. vol. 92, 1122–1136. doi: 10.1109/JPROC.2004.829009

Schwan, H. (1994). Electrical properties of tissues and cell suspensions: mechanisms and models. In Proceedings of 16th Annual International Conference of the IEEE Engineering in Medicine and Biology Society (IEEE), A70–A71. doi: 10.1109/IEMBS.1994.412155

Schwartz, Z., Simon, B. J., Duran, M. A., Barabino, G., Chaudhri, R., and Boyan, B. D. (2008). Pulsed electromagnetic fields enhance BMP-2 dependent osteoblastic differentiation of human mesenchymal stem cells. Journal of Orthopaedic Research 26, 1250–1255. doi: 10.1002/jor.20591

Sebastián, J. L., Muñoz San Martín, S., Sancho, M., and Miranda, J. M. (2004). Modelling the internal field distribution in human erythrocytes exposed to MW radiation. Bioelectrochemistry 64, 39–45. doi: 10.1016/j.bioelechem.2004.02.003

Smoluchowski, M. (1916). Drei Vortrage uber Diffusion, Brownsche Bewegung und Koagulation von Kolloidteilchen. Z. Phys., 557–585

Stewart, D. A., Gowrishankar, T. R., and Weaver, J. C. (2004). Transport lattice approach to describing cell electroporation: Use of a local asymptotic model. IEEE Transactions on Plasma Science 32, 1696–1708. doi: 10.1109/TPS.2004.832639

Sun, L.-Y., Hsieh, D.-K., Lin, P.-C., Chiu, H.-T., and Chiou, T.-W. (2009). Pulsed electromagnetic fields accelerate proliferation and osteogenic gene expression in human bone marrow mesenchymal stem cells during osteogenic differentiation. Bioelectromagnetics, n/a–n/a doi: 10.1002/bem.20550

Tandon, N., Goh, B., Marsano, A., Chao, P.-H., Montouri-Sorrentino, C., Gimble, J., et al. (2009). Alignment and elongation of human adipose-derived stem cells in response to direct-current electrical stimulation. In 2009 Annual International Conference of the IEEE Engineering in Medicine and Biology Society (IEEE), 6517–6521. doi: 10.1109/IEMBS.2009.5333142

Tonge, P. D., Olariu, V., Coca, D., Kadirkamanathan, V., Burrell, K. E., Billings, S. A., et al. (2010). Prepatterning in the stem cell compartment. PLoS ONE doi: 10.1371/journal.pone.0010901

Vajrala, V., Claycomb, J. R., Sanabria, H., and Miller, J. H. (2008). Effects of oscillatory electric fields on internal membranes: An analytical model. Biophysical Journal 94, 2043–2052. doi: 10.1529/biophysj.107.114611

Valic, B., Golzio, M., Pavlin, M., Schatz, A., Faurie, C., Gabriel, B., et al. (2003). Effect of electric field induced transmembrane potential on spheroidal cells: theory and experiment. Eur Biophys J 32, 519–528. doi: 10.1007/s00249-003-0296-9

Ying, W. and Henriquez, C. S. (2007). Hybrid finite element method for describing the electrical response of biological cells to applied fields. IEEE Trans Biomed Eng 54, 611–620

