## Supplement Information for "A theoretical framework to study the influence of electrical fields on mesenchymal stem cells"

### Supplementary Material

#### 1 DERIVATION PROCEDURE

Equations (5) and (10) in the main text are solved using Laplace technique to obtain equations for the total number of cells and the total ALP activity in the stimulation chamber. For the purpose of showing our solution approach, we combine the two equations into one equation, using the parameters choice given in Table (1) and (3), as follows,

$$\frac{\partial n(a,t)}{\partial t} \;=\; k_d n(a,t)\int_0^\infty \delta(a-a')da' + d_o\frac{\partial n(a,t)}{\partial a} - k_d n(a,t) \quad . \tag{S1}$$

The Laplace transform of $n(a,t)$ is defined as,

$$\tilde{n}(x,t) = \int_0^\infty e^{-ax} n(a,t)\, ds \quad . \tag{S2}$$

From Equation (S2) we obtain the following two quantities,

$$N \;=\; \tilde{n}(0,t), \tag{S3}$$

$$\Phi \;=\; -\frac{\partial \tilde{n}(x,t)}{\partial x}|_{x=0} = -\tilde{n}'(x,t)|_{x_0}, \tag{S4}$$

where, $N$ and $\Phi$ represent the total number of cells and the total ALP activity in these cells at time $t$, as defined in the main text. Using the definition given in Equations (S3) and (S4), the Laplace transformation of Equation (S1) is,

$$\frac{\partial \tilde{n}(x,t)}{\partial t} \;=\; k_d\tilde{n}(x,t) - d_o x\Big(\frac{\partial \tilde{n}(x,t)}{\partial x}\Big) - k_f\tilde{n}(x,t) \quad . \tag{S5}$$

where we have used the initial condition $n(a,t=0)=0$. Substituting $x=0$ in Equation (S5), we get, for the total number of ALP expressing cells,

$$\frac{dN}{dt} = k_d N - k_f N \tag{S6}$$

Differentiating Equation (S5) with respect to $x$ we get,

$$\frac{\partial \tilde{n}'(x,t)}{\partial t} \;=\; k_d\tilde{n}'(x,t) - d_o\Big(\frac{\partial \tilde{n}(x)}{\partial x}\Big) - d_o x\Big(\frac{\partial^2 \tilde{n}(x)}{\partial x^2}\Big) - k_f\tilde{n}'(x,t) \quad . \tag{S7}$$

Insert $x=0$ in Equation (S5) and using the identity given in Equation (S4), we get, for the total ALP activity in the cell culture chamber,

$$\frac{d\Phi}{dt} = k_d\Phi - d_o\Phi - k_f\Phi \quad . \tag{S8}$$
